# Context-Dependent Modulation of Astrocytic Ca^2+^ Signals by Mitochondria - A Computational Study

**DOI:** 10.1101/2025.07.14.664772

**Authors:** Thiago Ohno Bezerra, Antonio C. Roque

## Abstract

Mitochondria are one of the major regulators of intracellular Ca^2+^ in the cells, uptaking this ion through the mitochondrial Ca^2+^ uniporter (MCU) and releasing by the mitochondrial permeability transition pore (mPTP). Astrocytes respond to neurotransmitters and other stimuli by increasing the intracellular Ca^2+^ concentration, a process called ^2+^ signaling. However, it is not clear how mitochondria interact with the neurotransmitter-triggered Ca^2+^ responses in astrocytes. To explore this mechanisms, we expanded a previous compartmental model of astrocytes developed by our group including the mitochondrial MCU and mPTP mechanisms controlling the Ca^2+^ response. We simulate glutamatergic and dopaminergic inputs, modeled as Poisson processes, that promote the synthesis of IP_3_ through the PLC pathway. Here, we used a unipolar and a bifurcated-terminal morphology models and, with exception for the distal compartments, every other compartment have mitochondria. Simulations revealed that mitochondria modulate the Ca^2+^ response in a context-dependent manner. For weak glutamatergic input, they reduce the frequency of Ca^2+^ oscillations and the distance these signals propagate from the terminal regions. However, for strong glutamatergic input and in the presence of dopamine, mitochondria enhance the Ca^2+^ response by reducing the Ca^2+^-dependent IP_3_ degradation. Our findings provide computational evidence that mitochondria have a critical role in shaping the spatial organization of Ca^2+^ singling in astrocytes.

## 1 INTRODUCTION

Mitochondria are central for cell metabolism ^1^. Intramitochondrial Ca^2+^ is a major regulator of mitochondrial metabolism, being associated with the modulation of both oxidative phosphorylation and the synthesis of ATP ^1^. However, in addition to their metabolic function, mitochondria are also one of the major contributors to the Ca^2+^ activity in astrocytes ^2,3^. Although traditionally associated with support functions, as metabolic support to neurons, neurotransmitter homeostasis, and regulation of extracellular ions ^4^, recent reports showed that astrocytes are involved in the control of the excitatory-inhibitory balance ^5^, synaptic plasticity ^6^, neural oscillations ^7,8^, and processing of sensory stimuli ^9,10^, impacting long- and short-term memory, social behavior, and motor responses ^8,11,5^. Astrocytes express most of the neurotransmitter and neuromodulators receptors on their membrane, such as the metabotropic receptors for glutamate, α_1_ for noradrenaline, and D_1_/D_2_ for dopamine ^12,13,14,10,15^. These receptors activate the phospholipase-C (PLC) pathway, which promotes the synthesis of IP_3_ by a Ca^2+^ sensitive mechanism. IP_3_ interacts with IP_3_ channels on the ER membrane that releases Ca^2+^ to the cytosol. This is terminated by either IP_3_R channels closing or by IP_3_ degradation, which is dependent on intracellular Ca^2+^. Ionic channels, as NMDA receptors and transient receptor potential ion channels ^16,12,6,17^, and exchangers and transporters, as the Na^+^/Ca^2+^-exchanger and plasma membrane Ca^2+^ ATPase ^18,19,20^, also regulate intracellular Ca^2+^ in astrocytes. This increase in intracellular Ca^2+^ is often called Ca^2+^ signals, oscillations or microdomains, depending on time, frequency and location. These Ca^2+^ responses can be further classified as spontaneous or elicited responses ^17^, whether triggered by neurotransmitter, and into local or global responses ^21,17,15^, whether it takes the whole cell. Global elicited responses were associated with neuromodulators, as noradrenaline and dopamine ^14,15^, while local responses to synaptic glutamate, mitochondria, ionic channels and transporter on the plasmatic membrane ^16,19,17,15,12^. Ca^2+^ signals in astrocytes can promote the release of the so called gliotransmitters, such as glutamate, ATP and D-serine ^22,23,17^. So, studying the Ca^2+^ response in astrocytes is essential to understand how these cells can modulate neural activity.

In addition to the mechanisms mentioned above, mitochondria have also been associated with spontaneous and elicited Ca^2+^ responses in astrocytes ^2,3,24^. The mitochondrial Ca^2+^ uniporter (MCU) ^3,25,24^ uptakes the Ca^2+^ from the cytosol into the mitochondria, increasing the intra-mitochondrial Ca^2+^ up to tens of µM ^25^. This uniporter is regulated by the mitochondria electric potential and redox state of the cell ^26,27^. On the other hand, Ca^2+^ is released through the Na^+^/Ca^2+^ exchanger (NCX) and the mitochondrial permeability transition pore (mPTP) ^2,24^. Although mitochondrial Ca^2+^ overload and redox imbalance were often associated with prolonged opening of mPTP ^24^, a process that can lead to cell death, recent studies have shown that transient opening of mPTP can occur under physiological condition and regulate cell homeostasis ^28^. In astrocytes, some studies have associated both MCU and mPTP to Ca^2+^ signaling ^2,3^. Since the mitochondria can also be found closely apposed to the ER ^25,3,24^, they are particularly suited as one of the major regulators of the Ca^2+^ response. As some astrocyte regions have smaller ER compartments or even are devoid of them ^19,29,30,24^, mitochondria that are present in these regions can also contribute to Ca^2+^ signaling ^30,31^. Interestingly, mitochondria are mobile organelles that in astrocytes move towards regions with higher Ca^2+^ activity and glutamate uptake ^32,24^. In this context, functional alterations in both astrocytes and mitochondria have been associated with aging ^33^, neurodegenerative diseases, such as Alzheimer’s and Parkinson’s diseases ^34,35^, and psychiatric disorders, such as schizophrenia ^36,37,38^. In particular, alterations in the expression of some of the mitochondrial enzymes that are regulated by Ca^2+^ were associated with schizophrenia ^39^. So, instead of just controlling the cell metabolism, mitochondria are important regulators of astrocyte Ca^2+^ responses, and possibly implicated in the modulation of neural activity and plasticity through the astrocyte.

As mentioned above, Ca^2+^ response in astrocytes involves complex biochemical pathways ^12^ and some studies using mathematical and computational modeling were develop to account for this response. These models can be classified into three categories: a) De Young and Keizer (1992) and Li and Rinzel (1994) -type models; b) Höfer et al. (2002) -type models; and c) mixed type models ^40^. In the first case, they model the generation of Ca^2+^ signals considering the release of Ca^2+^ from intra-ER stores by the activation of IP_3_ channels. On the other hand, although simulating the same ER mechanism, those classified as Höfer type are reaction-diffusion models, detailing how spatial variables can influence astrocytic Ca^2+^ response. But, among these models, only two have addressed the influence of mitochondria on astrocyte Ca^2+^ signaling. Komin and colleagues ^18^ modeled the mitochondria Ca^2+^ release, but they did not specify the mechanism responsible for the Ca^2+^ release, implemented a Ca^2+^ influx into mitochondria nor tested directly the effect of this mechanism on the astrocyte response. In contrast, Diekman *et al*. ^41^, extending other models by Magnus and Keizer ^42^, and Oster *et al*. ^43^, developed a computational model to investigate the neuroprotective role of mitochondrial metabolism during stroke. In this model they included the mitochondrial Ca^2+^ influx through the MCU and efflux through the NCX ^41^. However, the mitochondria Ca^2+^ handling was not the main focus of this study. So, although implementing mitochondria Ca^2+^ mechanisms, the previous computational models did not investigate the impact of mitochondria on the Ca^2+^ response, less so how neurotransmitter elicited response could interact with mitochondria neither how the astrocyte morphology would impact those factors ^15^.

Here we present a compartmental astrocyte model developed as an extension of a previous computational model by our group ^15^ including both MCU and mPTP mitochondrial mechanisms regulating the intracellular Ca^2+^. With this model we explore how the glutamate and dopamine elicited responses interact with these mitochondrial mechanisms, showing that the mitochondria regulate astrocyte Ca^2+^ response in a context-dependent manner, inhibiting the response for weak glutamatergic stimulation but enhancing it for strong glutamatergic input or in the presence of dopamine. This process is mediated by the reduction of Ca^2+^-dependent IP_3_ degradation. Finally, we investigate how mitochondria gate the transmission of Ca^2+^ signals from the distal regions of an astrocytic process.

## 2 MODEL DESCRIPTION AND METHODS

The present model is an extension of a previous compartmental model of astrocytes developed by our group ^15^ and based on previous computational models of Ca^2+^ signaling in astrocytes and mitochondrial Ca ^2+^ handling ^44,45,46,47,19,48^. In the first subsection, we describe a single compartment with the variables and currents associated with the mitochondria mechanism. Next, we describe how the compartments are connected, the morphological models used in this study and the mitochondria

### 2.1 Single Compartment Model

Each compartment is subdivided into intra-ER, intramitochondrial, intracellular, and extracellular spaces (Fig. 1**A**) modeled as well mixed solutions and connected by current densities between subspaces. We modeled the following variables: intracellular and extracellular Ca^2+^ concentrations ([Ca^2+^]_i_ and [Ca^2+^]_e_, respectively); ER Ca^2+^ concentration ([Ca^2+^]_ER_); mitochondria Ca^2+^ concentration ([Ca^2+^]_m_), intracellular IP_3_ concentration ([IP_3_]); the fraction of open IP_3_ channels (*h*); intracellular and extracellular Na^+^ concentrations ([Na^+^]_i_ and [Na^+^]_e_, respectively); intracellular and extracellular K^+^ concentrations ([K^+^]_i_ and [K^+^]_e_, respectively); the compartment potential (*v*); and extracellular glutamate ([Glu]) and dopamine ([DA]) concentrations. With exception for the rate of change of intracellular and intramitochondrial Ca^2+^ concentrations and the currents connecting these subspaces – which we describe here in the main text –, the expressions for the rate of change of the remaining variables, the current densities, and model parameters are the same as in the previous computational model ^15^.

**FIGURE 1.**
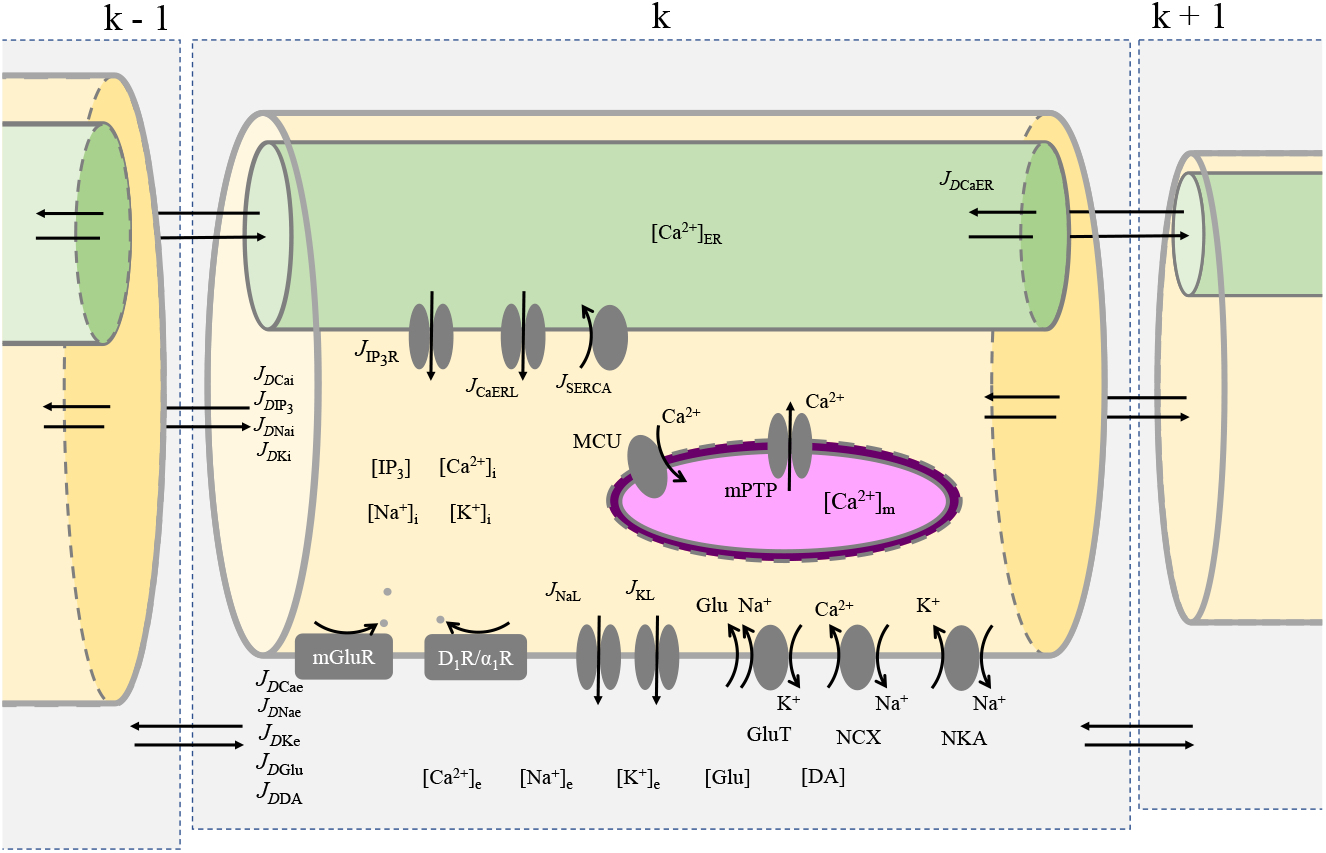
Schematic representation of compartment variables and currents. Each compartment is subdivided into extracellular (gray), intra-mitochondria (purple), intracellular (yellow) and intra-ER (green) spaces. The intracellular Ca^2+^ concentration ([Ca^2+^]_i_) is regulated by the release from the ER through IP_3_R channels 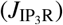, leak from ER (*J*_CaERL_), SERCA pump uptake (*J*_SERCA_), uptake through the MCU (*J*_MCU_), release through the mPTP (*J*_mPTP_), the NCX (*J*_NCX_), and diffusion between compartments (*J*_*D*Cai_). Extracellular Ca^2+^ ([Ca^2+^]_e_) is controlled by the NCX and diffusion (*J*_*D*Cae_). ER subcompartments are also connected by Ca^2+^ diffusion currents (*J*_*D*CaER_). Intra-mitochondria Ca^2+^ is regulated by the MCU and mPTP. Intracellular IP_3_ concentration ([IP_3_]) is controlled by the PLCβ activated through the metabotropic glutamatergic (mGluR) and dopaminergic/noradrenergic receptors (D_1_R/α_1_R), the PLC*δ*, by the enzymes IP_3_-3K and IP-5P, and by diffusion between compartments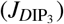. The intracellular ([Na^+^]_i_ and [K^+^]_i_) and extracellular concentrations ([Na^+^]_e_ and [K^+^]_e_) of the other ionic species (Na^+^ and K^+^) are regulated by the leak currents (*J*_NaL_ and *J*_KL_), glutamate transporter (*J*_GluT_), NCX, NKA (*J*_NKA_) and diffusion between compartments (*J*_*D*Nai_, *J*_*D*Nae_, *J*_*D*Ki_, and *J*_*D*Ke_). Extracellular glutamate ([Glu]) and dopamine ([DA]) concentrations determined by the presynaptic release events, simulated as Poisson processes, and by diffusion between extracellular subcompartments (*J*_*D*Glu_ and *J*_*D*DA_) distribution within the astrocyte. The remaining variables and currents implemented in this model, the compartment parameters, and the stimulation protocols are given in Text S1.

The intracellular and mitochondrial Ca^2+^ concentrations are given by:

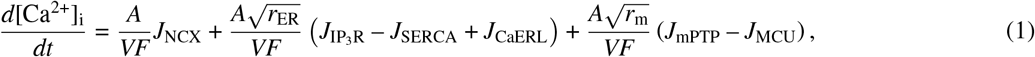

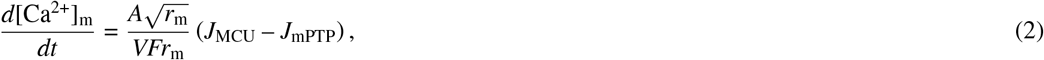

where *A* is the superficial area of the compartment, *V* the volume of the compartment, *F* the Faraday’s constant, *r*_ER_ the ratio between the ER and cytosol volumes (Eq. 5), *J*_NCX_ the plasma membrane current though the NCX, 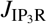 the current through the IP_3_ receptor channels on the ER membrane (Eq. S12), *J*_SERCA_ the current through the SERCA transporter (Eq. S13), *J*_CaERL_ is the leak current density through the ER (Eq. S14), *r*_m_ is the ratio between mitochondria and cytosol volumes, *J*_mPTP_ is the efflux current density through the mPTP (Eq. 4), and *J*_MCU_ is the Ca^2+^ uptake current by the MCU into the mitochondria (Eq. 3).

The MCU and mPTP current densities are given by:

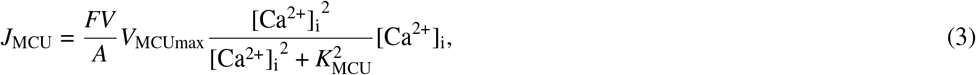

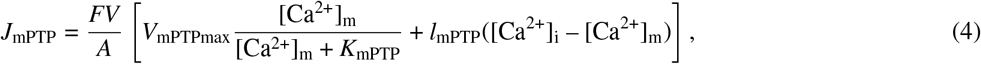

where *V*_MCUmax_ is the maximum transport rate of the MCU, *K*_MCU_ is the Ca^2+^ affinity constant to the MCU, *V*_mPTPmax_ is the maximum efflux rate of Ca^2+^ through the mPTP, *K*_mPTP_ is the Ca^2+^ affinity to mPTP and is *l*_mPTP_ the passive Ca^2+^ efflux rate through the mPTP. This mitochondrial model is based on a previous model by Pivovarova *et al*. ^45^, and Friel and Tsien ^44^. The parameters of these currents were adapted to the astrocyte model as described in the section Mitochondria Model in Text S1.

To account for the size difference of the ER and the mitochondria according to their position within the astrocyte ^29,30,49,31^, we implemented the factor *r*_ER_ and *r*_m_ representing, respectively, the volume fraction of the ER and mitochondria to the cytosol. *r*_ER_ is given by:

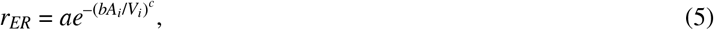

where the parameters *a* = 0.15, *b* = 0.073 µm and *c* = 2.34 were fitted using experimental data ^29^, and *A*_*i*_/*V*_*i*_ is surface-volume ratio of the compartment.

As a simplification, we modeled the total volume of mitochondria in one compartment as a cylinder with constant length *l* = 1 µm, same length of each compartment, and radius *ϕ*_*m*_ as function of the compartment radius *ϕ*:

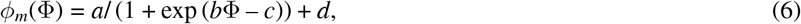

where *a, b, c* and *d* are free parameters fitted using the data of mitochondria thickness from adult mouse somatosensory cortical astrocytes ^49^, and Φ the compartment radius. So, the ratio between mitochondria (*V*_*m*_) and cytosol (*V*_*i*_) volumes is given by *r*_m_ = *V*_*m*_/*V*_*i*_. More details on this parameter fitting are given in the Text S1.

### 2.2 Compartment Coupling and Cell Morphologies

Compartments are connected by diffusive fluxes of Ca^2+^, Na^+^, K^+^, IP_3_, glutamate and dopamine between intracellular, extracellular, and ER subspaces of consecutive compartments. Here we implemented the mitochondria as isolated compartments, so we did not consider here the mitochondrial complexes that are reported in some studies ^31^. The generic expression for the rate of change of the concentration of ion/molecule X in compartment *j* due to the diffusive currents from/to a neighboring compartment *k* is given by ^50^:

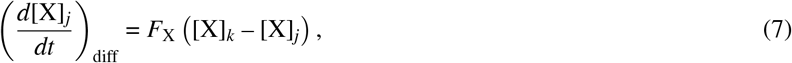

where *F*_X_ (s^-1^) is the coupling strength between compartments *j* and *k* for the ion/molecule X. The coupling strength depends on the geometry of the compartments and on the diffusion coefficient *D*_X_ ^15^. More details regarding the diffusive fluxes and the related parameters are given in the Text S1.

We first simulated the model with a unipolar morphology composed of a somatic spherical compartment followed by 20 cylindrical compartments (Table S1 in Text S1), representing a single process in an astrocyte ^15^. In the last test, we used a bifurcated-terminal morphology (24 compartments, Fig. 8**A** and **B**, and Table S2 in Text S1), composed of a somatic spherical compartment from which a linear process emanates that bifurcates close to the distal region. Unless otherwise stated, since mitochondria are not present everywhere within the astrocyte ^49,31^, in the present model the distance between compartments with mitochondria is two, so starting from the soma, the mitochondria are present in every other compartment. Distal compartments with radii below 0.2 µM also do not have mitochondria, since mitochondria are thicker than these compartments. We tested two bifurcated-terminal morphologies, one with mitochondria at the bifurcation point and the other without. More details on the morphological models used here are given in Text S1.

### 2.3 Model Input and Output

In the present study, we simulate both glutamatergic and dopaminergic transmission to astrocytes. Both stimuli trigger Ca^2+^ responses by activating the PLC pathway, promoting IP_3_ synthesis and release of Ca^2+^ from the ER ^51,52,53,15^. Glutamate also activates the glutamate transporter (GluT), inducing Ca^2+^ uptake from extracellular space through the NCX by altering the Na^+^ gradient across the astrocyte membrane (Fig. 1) ^12,19,54,15^. Both stimuli were simulated as Poisson processes with frequencies *v*_*g*_, for glutamate, and *v*_*d*_ for dopamine, given in Hz and held constant during each simulation trial. To indicate the compartments that are under stimulation, we use the following notation: CX-Y, where C stands for compartment, X for the first compartment under stimulation, and Y the last compartment. With exception for the test in which the position of glutamatergic input and mitochondria were changed (Figures S10 and S12), glutamate always stimulated the distal compartments (C18-20), simulating localized inputs from tripartite synapses ^54^, while dopamine stimulated the entire cell (C1-20), simulating the volume transmission characteristic of neuromodulators ^55^.

Ca^2+^ response is analyzed with respect to the frequency (*f*_Ca2+_) and location of the Ca^2+^ oscillations. Ca^2+^ signals are counted every time the intracellular Ca^2+^ concentration crosses the threshold [Ca^2+^]_th_ = 015 µM ^48,15^. The frequency of Ca^2+^ oscillations is defined here as the number of Ca^2+^ signals triggered during the stimulation period divided by the trial duration (*t* = 100 s). Here we also analyzed the distance Ca^2+^ signals propagate from the terminal regions, counted as the number of compartments activated counting from the last compartment.

## 3 RESULTS

First, to compare the model response with and without the mitochondrial mechanisms, we simulated the unipolar model of astrocyte with glutamatergic (*v*_*g*_ = 5 Hz, C18-20) and dopaminergic (*v*_*d*_ = 1 Hz, C1-20) inputs. Here, we focused our analysis on compartments C13-16. Details regarding the response of the remaining compartments (FigS2) and the simulations with only one neurotransmitter (FigS3) are given in the Text S1.

Glutamatergic and dopaminergic stimulation trigger Ca^2+^ signals in all compartments in both models (Fig. 2 and FigS2**A**). Compartments with mitochondria have Ca^2+^ signals with smaller amplitudes. However, the frequencies of Ca^2+^ oscillations 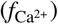are higher in all compartments of the model with mitochondria. The intracellular Ca^2+^ oscillation is followed by an increase in the mitochondrial Ca^2+^ (Fig. 2**B**). Interestingly, the IP_3_ concentration of all compartments is higher in the mitochondria model, so mitochondria prevented the degradation of IP_3_ during a Ca^2+^ signal (Fig. 2**C**). This is also evidenced in the Ca^2+^-IP_3_ space by the overall smaller Ca^2+^ oscillation in the compartments with mitochondria but higher IP_3_ (FigS2**B**). Similar to the stimulation with both glutamate and dopamine, the mitochondria also enhance the response to either glutamate input alone (*v*_*g*_ = 5 Hz, FigS3**A**) or dopaminergic input alone (*v*_*d*_ = 1 Hz, FigS3.**B**). These results suggest that the mitochondria enhance the astrocyte response by either temporally reducing the intracellular Ca^2+^, and/or increasing the intracellular IP_3_ concentration.

**FIGURE 2.**
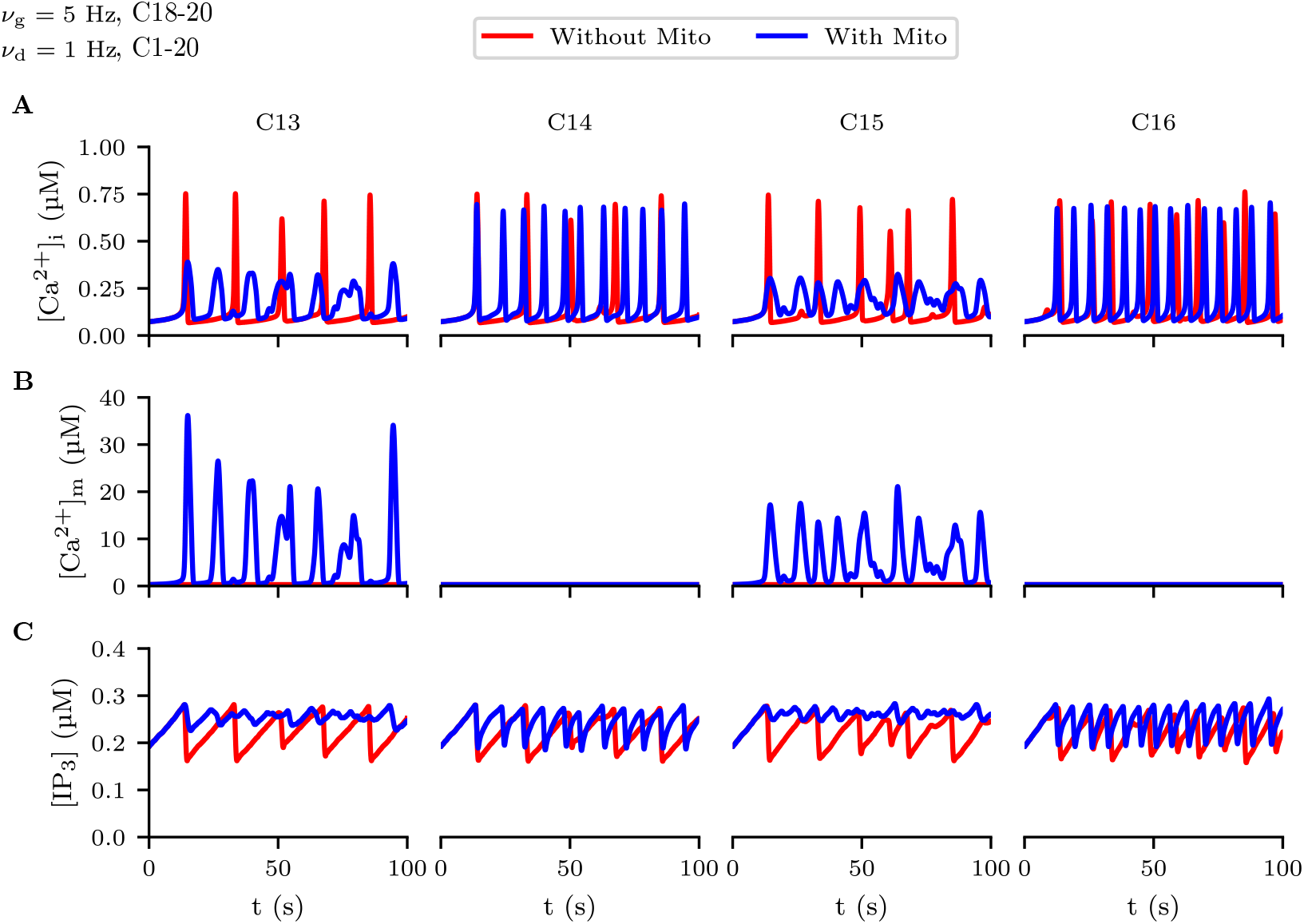
Glutamatergic and dopaminergic stimulation trigger Ca^2+^ signals in the astrocyte models. Compartments C13 and C15 have mitochondria. **A**. Intracellular Ca^2+^ concentration. **B**. Intramitochondrial Ca^2+^ concentration. **C**. Intracellular IP_3_ concentration.

**FIGURE 3.**
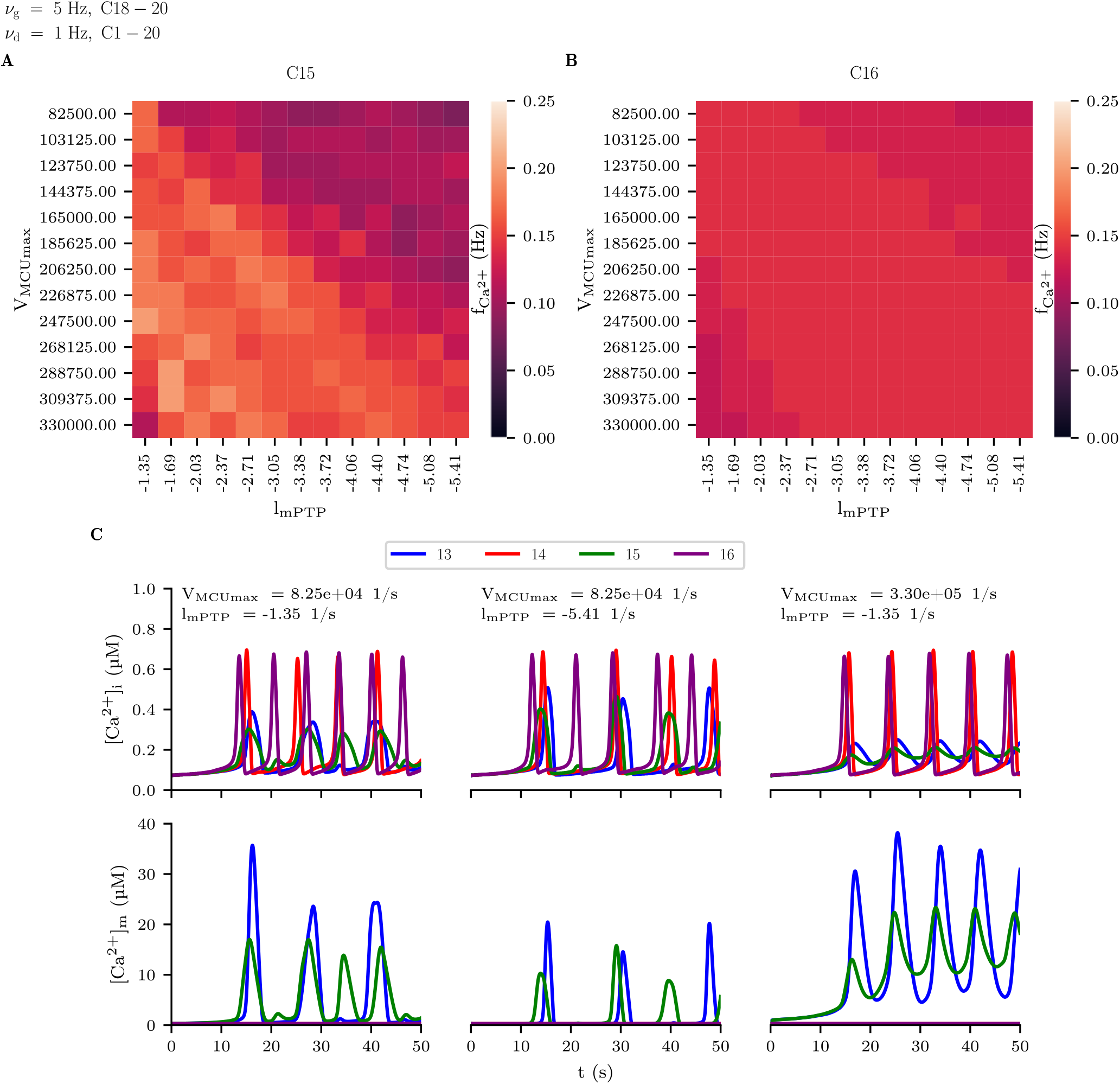
Mitochondria Ca^2+^ uptake and release rates affect the frequency and pattern of Ca^2+^ oscillations. 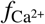in compartment C15 (**A**) and C16 (**B**). **C** Intracellular ([Ca^2+^]_*i*_) and intramitochondrial ([Ca^2+^]_*m*_) Ca^2+^ traces for three combinations of MCU uptake (V_MCUmax_) and mPTP release (l_mPTP_) rates.

To test the effect of mitochondrial Ca^2+^ uptake through the MCU and release by the mPTP on Ca^2+^ oscillations, we simulated the astrocyte model using different rates of maximum MCU uptake (*V*_MCUmax_, Eq. 3) and mPTP release (*l*_mPTP_, Eq. 4). Increasing the MCU Ca^2+^ uptake can decrease or increases the 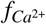in the compartments with mitochondria depending on the *l*_mPTP_ (Fig. 5**A** and FigS4.**A**). With low mPTP Ca^2+^ release, increased uptake reduces the frequency of response. In contrast, for higher release rates, increasing the MCU uptake also increases the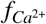. Changing the uptake and release has a minor effect in compartments without mitochondria (Fig. 5**B** and FigS4.**A**). So, the effect of mitochondrial Ca^2+^ handling depends on the *V*_MCUmax_/*l*_mPTP_ ratio, with the highest 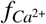achieved when there is a balance between uptake and release of Ca^2+^ by mitochondria (FigS4**B**).

Different patterns of Ca^2+^ oscillations are observed for different combinations of *V*_MCUmax_ and *l*_mPTP_ (Fig. 5**C**). Smaller MCU uptake and mPTP release produce smaller amplitudes of intracellular Ca^2+^ oscillations in compartments with mitochondria, which is associated with higher 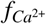. The amplitude of intramitochondrial Ca^2+^ oscillations are also higher in this case. The amplitude of intracellular Ca^2+^ oscillations increases with the same MCU uptake but higher mPTP release, but the 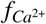is reduced. The amplitude of intramitochondrial Ca^2+^ oscillations is also smaller, showing isolated Ca^2+^ peaks (Fig. 5**C**). Finally, higher MCU uptake with lower mPTP release rates leads to stable intracellular Ca^2+^ oscillations and sustained increase in the intramitochondrial Ca^2+^ (Fig. 5**C**).

Together, these results show that both mitochondrial Ca^2+^ uptake and release control the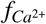, with the highest 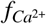obtained for balanced MCU uptake and mPTP release, which is associated with smaller amplitudes of Ca^2+^ oscillations. Different patterns of intracellular and mitochondrial Ca^2+^ oscillations can also be triggered by changing the combinations of *V*_MCUmax_ and *l*_mPTP_.

### 3.1 IP_3_-Ca^2+^ Interaction

As shown before, higher 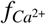induced by mitochondria is associated with smaller amplitude of intracellular Ca^2+^ oscillations and with higher intracellular IP_3_. To explore the mechanism by which mitochondria enhance the Ca^2+^ response, we simulated the model changing the strength of the Ca^2+^-IP_3_ interaction. In this test, we changed the parameters associated with the Ca^2+^-dependent IP_3_ degradation (K_D_, S23), the Ca^2+^-dependent inhibition of PLC (K_p_, S21), the affinity coefficient of Ca^2+^ to PLC (K_*π*_, S21), and the Ca^2+^-independent IP_3_ degradation rate (r_5P_, S24). In this test, we analyzed the difference in 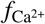between the model with mitochondria to the model without mitochondria, with zero difference indicating a similar response between the models. Here we also stimulated the unipolar model with glutamate (*v*_*g*_ = 5 Hz, C18-20) and dopamine (*v*_*d*_ = 1 Hz, C1-20). Complementary to the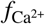, we also analyzed the time average Ca^2+^ efflux through the IP_3_ channel (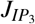, S12, FigS6), and the time average of IP_3_ degradation by IP3-3K (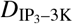, S23, FigS7). In the main text we provide the results for the 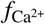changing the *K*_*D*_, the remaining results are given in the Text S1.

Only increasing the Ca^2+^-dependent IP_3_ degradation reduces the difference in 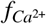between the models (Fig. 4**A** and FigS5). Decreasing the K_D_, which reduces the Ca^2+^ release from ER and increases the degradation of IP_3_ (FigS6 and FigS7, Eq. S23), also reduces the 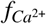in both models (Fig. 4**A**). In contrast, there was no difference in the model responses by changing K_p_ and K_*π*_ in any of the three metrics considered (FigsS5, FigsS6 and FigsS7). Similar to the Ca^2+^-dependent degradation, increasing the *r*_5*P*_ also reduced the 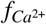(FiS5) and the Ca^2+^ efflux from ER (FigS6), with no major effect over the Ca^2+^-dependent degradation (FigS7). So, increasing the IP_3_ degradation reduce the difference between the models.

**FIGURE 4.**
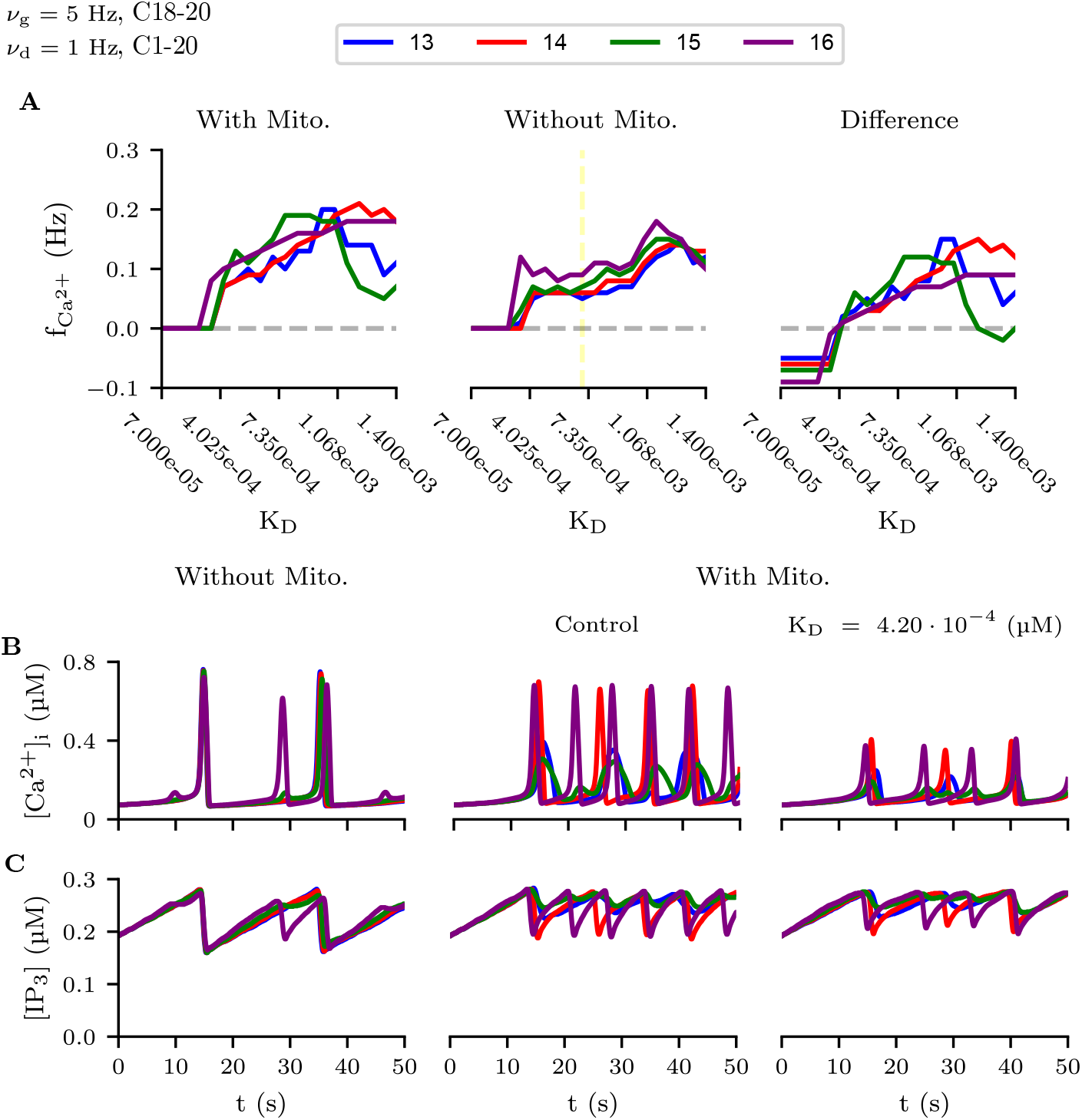
Mitochondria reduce Ca^2+^-dependent IP_3_ degradation. *K*_*D*_ controls the Ca^2+^ affinity to the IP_3_-3K. **A** Frequency of Ca^2+^ oscillations for different *K*_*D*_. The yellow line indicates the reference to calculate the difference between the model without mitochondria with the same parameter value. Intracellular Ca^2+^ (**B**) and intracellular IP_3_ concentrations (**C**) of the models without mitochondria, the model with mitochondria and default *K*_*D*_, and the model with mitochondria but reduced *K*_*D*_.

Decreasing the *K*_*D*_ reduces the 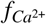in the model with mitochondria to a frequency similar to the model without mitochondria, but reduces the amplitude of the Ca^2+^ signals in all compartments (Fig. 4**B**). Similarly, the IP_3_ concentration is also reduced in the model with mitochondria and smaller *K*_*D*_ (Fig. 4**C**). However, the IP_3_ dynamics of the model without mitochondria is not fully recovered by changing the *K*_*D*_ in the model with mitochondria, as the amplitude of IP_3_ oscillation is still different between the models. So. although increasing the Ca^2+^-dependent IP_3_ degradation produces a response in the model with mitochondria that is similar to the model without mitochondria, the amplitude of Ca^2+^ and IP_3_ are still different.

To further test the hypothesis that mitochondria enhance the astrocyte response by reducing the Ca^2+^-dependent IP_3_ degradation, we simulated a trial with reduced MCU uptake rate while decreasing the IP_3_ degradation. The same glutamatergic (*v*_*g*_ = 5 Hz, C18-20) and dopaminergic (*v*_*d*_ = 1 Hz, C1-20) inputs are applied here. Here we show the results for V_MCUmax_ = 41.25 · 10^3^ s^−1^ and K_D_ = 0.84 µM. The 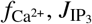 and 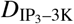for a range of *K*_*D*_ are given in the Text S1 (FigS8).

Inhibition of MCU uptake reduces the intracellular and intramitochondrial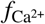, increases the amplitude of intracellular Ca^2+^ oscillations and triggers isolated intramitochondrial Ca^2+^ peaks (Fig. 5). However, decreasing the Ca^2+^-dependent degradation by increasing *K*_*D*_ recovers the reduction in the intracellular 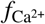caused by MCU inhibition and further increases the amplitude of intracellular Ca^2+^ oscillations. The amplitude of the intramitochondrial Ca^2+^ oscillations increases with the reduction of IP_3_ degradation, but the amplitude is still below the control condition and the stimulation triggers isolated Ca^2+^ peaks. So, the effects of inhibiting the MCU uptake on the intracellular and intramitochondrial Ca^2+^ responses are attenuated by reducing the Ca^2+^-dependent IP_3_ degradation.

**FIGURE 5.**
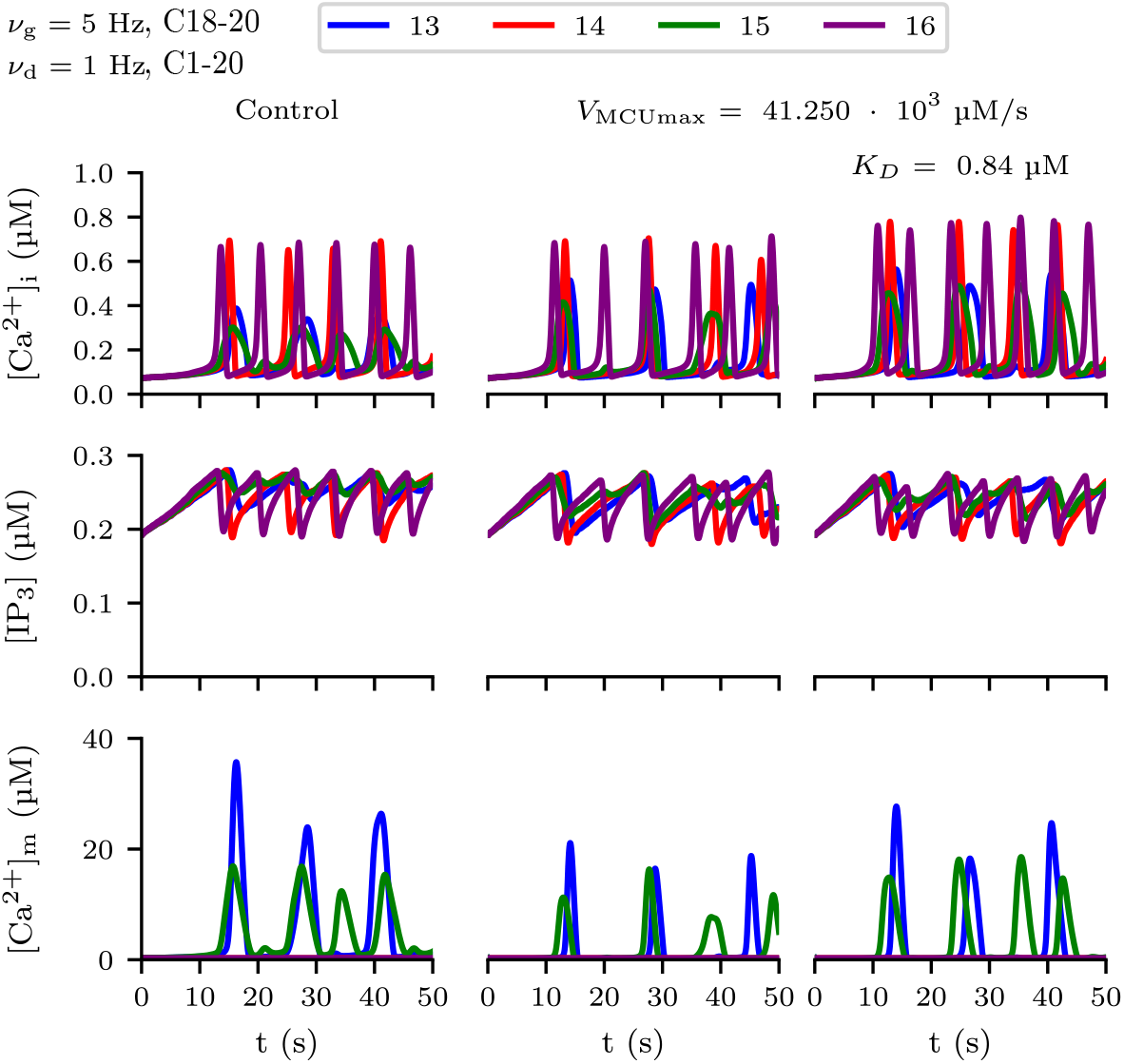
Reducing IP_3_ degradation attenuates the reduction of MCU Ca^2+^ uptake by mitochondria on the intracellular Ca^2+^ oscillation. In this test, the second and third columns show the traces for *V*_MCUmax_ = 41.25 10^3^ µM /s; and the third column with *K*_*D*_ = 0.84 µM. Distal compartments are simulated with glutamate (*v*_*g*_ = 5 Hz) and all compartments with dopamine (*v*_*d*_ = 1 Hz).

Together, these results suggest that mitochondria affect the astrocyte response by temporally reducing intracellular Ca^2+^ concentration which reduces the Ca^2+^-dependent IP_3_ degradation, allowing enough IP_3_ build-up to activate the IP_3_R and so increasing the Ca^2+^ response. Although mitochondria are present only in some compartments of the model, these organelles modulate the response in all compartments.

### 3.2 Mitochondria enhances the propagation of Ca^2+^ signals

As shown before, mitochondria enhance the Ca^2+^ response to glutamate and dopamine by preventing the IP_3_ degradation. In a previous paper by our group, we showed that dopamine enhances the glutamate-triggered responses and improves the propagation distance of Ca^2+^ signals from the distal compartments, facilitating the communication between different branchlets in the astrocyte model ^15^. To test whether mitochondria could also affect the propagation distance of Ca^2+^ signals in the astrocyte model, we simulated the unipolar astrocyte model (Fig. 6**A**) with different frequencies of dopaminergic (*v*_*d*_ = 0, 0.005, 0.01 and 0.05 Hz, C1-20) and glutamatergic (*v*_*g*_ from 0 to 10 Hz, C18-20) inputs and analyzed the propagation distance, which is defined here as the number of activated compartments counting from the most distal one. More details on the stimulation protocols are given in the Text S1.

**FIGURE 6.**
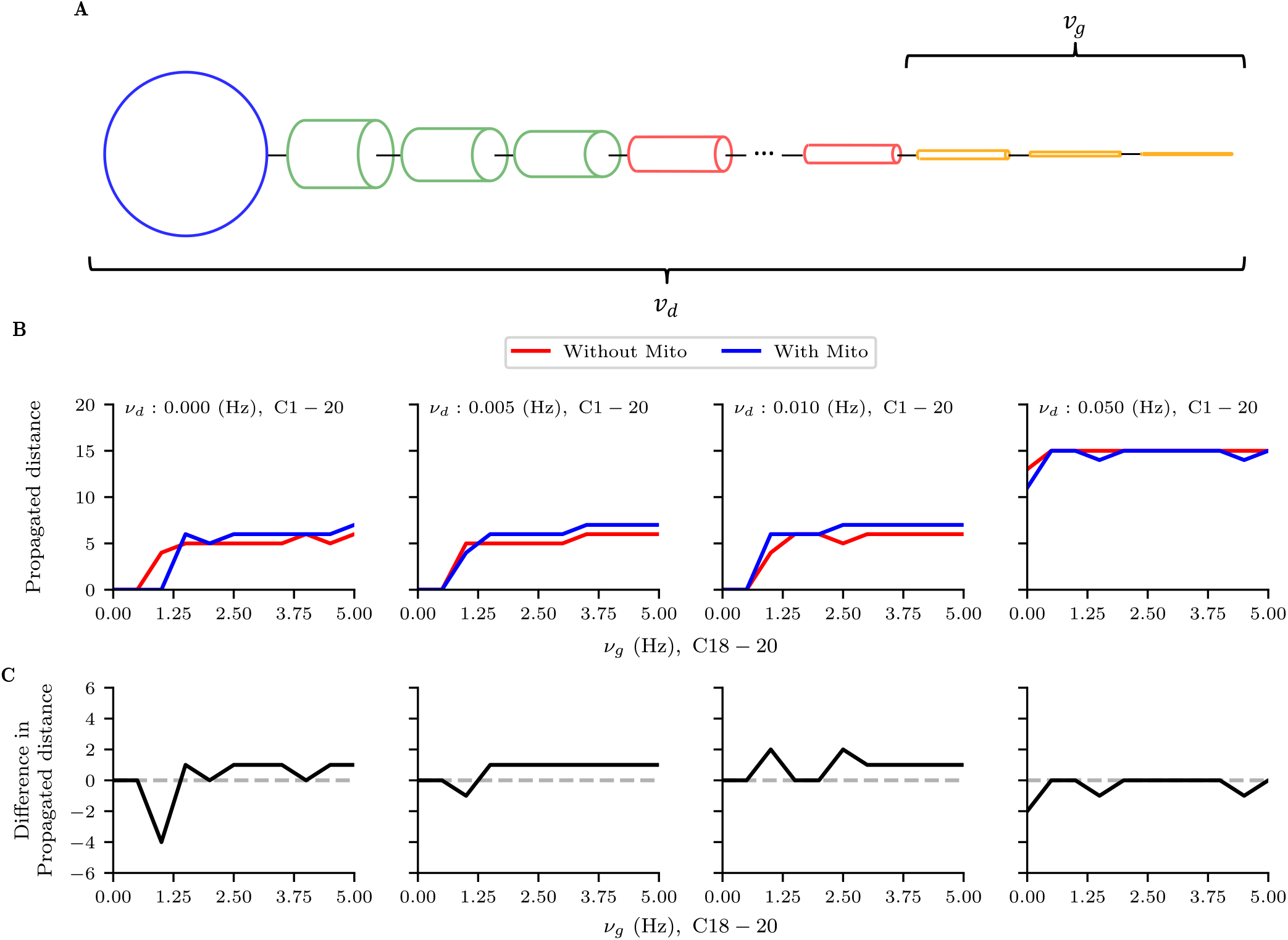
Mitochondria gates the transmission of Ca^2+^ signals in the linear astrocytic branch. The distal compartments are stimulated with glutamate (*v*_*g*_ from 0 to 5 Hz) while the whole cell is stimulated with dopamine (*v*_*d*_ = 0, 0.005, 0.01 or 0.05 Hz). **A** Schematic representation of the model morphology with the location of both inputs. **B** Propagated distance. **D** Difference in propagated distance between the models with and without mitochondria..

Without dopaminergic stimulation, mitochondrial have a bimodal effect on the propagation distance of Ca^2+^ signals triggered by glutamate. Mitochondria decrease the distance of propagation for low glutamatergic input (*v*_*g*_ < 1.5 Hz) but increase for higher frequencies (Fig. 6**B**). Taking the difference between the model with mitochondria and the model without mitochondria, the mitochondria reduce in 4 compartments the propagated distance of Ca^2+^ signals triggered by glutamate (*v*_*g*_ = 1 Hz, Fig. 6**C**). For*v*_*g*_ ≥ 1.5 Hz, the mitochondria increase the propagated distance up to one compartment. Low frequency dopaminergic input (*v*_*d*_ = 0.005 Hz) attenuates the mitochondrial reduction on the propagated distance (Fig. 6**C**). With*v*_*d*_ = 0.010 Hz mitochondria increase the propagation distance for all frequencies of glutamatergic stimulation (Fig. 6**C**). Finally, for the strongest dopaminergic input used in this test (*v*_*d*_ = 0.05 Hz) mitochondria had a small effect on the propagation distance, decreasing it in one compartment for some glutamatergic input frequencies (Fig. 6**C**). As higher dopaminergic input already triggers Ca^2+^ signals in almost all compartments, there is a minor effect of mitochondria on the propagated distance with stronger dopaminergic inputs. So, the mitochondria gates the transmission of Ca^2+^ from the distal regions in the absence of dopaminergic input and with weak glutamatergic stimulation, preventing that weaker inputs activate larger regions of the astrocyte. Dopamine reduces this glutamatergic threshold, allowing weaker stimuli to trigger responses that propagate along the astrocyte. This can be seem as a shift to the left in the propagated distance curve (Fig. 6**B**).

Intracellular Ca^2+^ also affect the mitochondria metabolism ^1^ and since the SERCA pump is energy-dependent, mitochondria could affect the propagation of Ca^2+^ by modulating Ca^2+^ uptake through SERCA pump. As a control test, we also simulated the model changing the maximum rate of SERCA pump (*v*_ER_, S14), indirectly simulating the effect of energy metabolism on Ca^2+^ handling. However, increasing the pumping rate reduces the propagation distance (FigS.8), the opposite we detected by comparing the models with and without mitochondria (Fig. 6**B**). To further test the mitochondrial gating effect, we also simulated a unipolar morphology in which only a single compartment has mitochondria and is stimulated in only one compartment. In this test we changed both the position of the mitochondria and the glutamatergic input (Fig.S9) and defined the propagation distance as the number of different compartments activated by the stimulation. Mitochondria closer to the distal compartments prevent the propagation of Ca^2+^ signals triggered in these compartments (Fig.S9). Stimulating the transition between the distal compartments and the intermediate compartments (C15-17), the mitochondria increases the propagation distance (Fig.S9). In the remaining compartments of the main branch, the mitochondria only prevented the propagation if these organelles were positioned close to the stimulation point. Finally, Ca^2+^ signals triggered by stimulating the somatic or proximal compartments do not propagate along the astrocyte (Fig.S9).

A similar effect to the propagation distance is observed by analyzing the 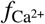(Fig. 7). Here we focused in the compartments C15-17, the data for the remaining compartments are given in the supplementary materials (FigS11). Compartments C15 and C17 have mitochondria. In compartments C16 and C17, mitochondria reduce the 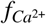for weak glutamatergic inputs (*v*_*g*_ ≤ 2 Hz) in the absence of dopaminergic transmission (Fig. 7). For higher input frequencies (*v*_*g*_ > 2 Hz), mitochondria increase 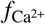(Fig. 7). In contrast with C16 and C17, mitochondria enhance the response of compartment C15 for any*v*_*g*_ (Fig. 7). Dopaminergic stimulation shifts the 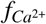curve to the left, reducing the minimum*v*_*g*_ needed for the mitochondria to enhance the compartment response to glutamatergic stimulation (Fig. 7). With the highest dopaminergic input simulated here (*v*_*d*_ = 0.05 Hz), the model with mitochondria has higher 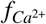for any*v*_*g*_ applied (Fig. 7). In both models, higher*v*_*g*_ triggers responses with higher 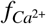in all compartment and for any*v*_*d*_ (Fig. 7). Similar to the propagation distance, in the unipolar morphology with one single mitochondria, the 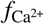of different compartments are also affected by the location of the mitochondria and the glutamatergic input (FigS12), in which co-localization in compartments C17 and C18 increased the response in these compartments, while the mitochondria closer to the distal regions decrease the response of compartments C19 and C20 (FigS12).

**FIGURE 7.**
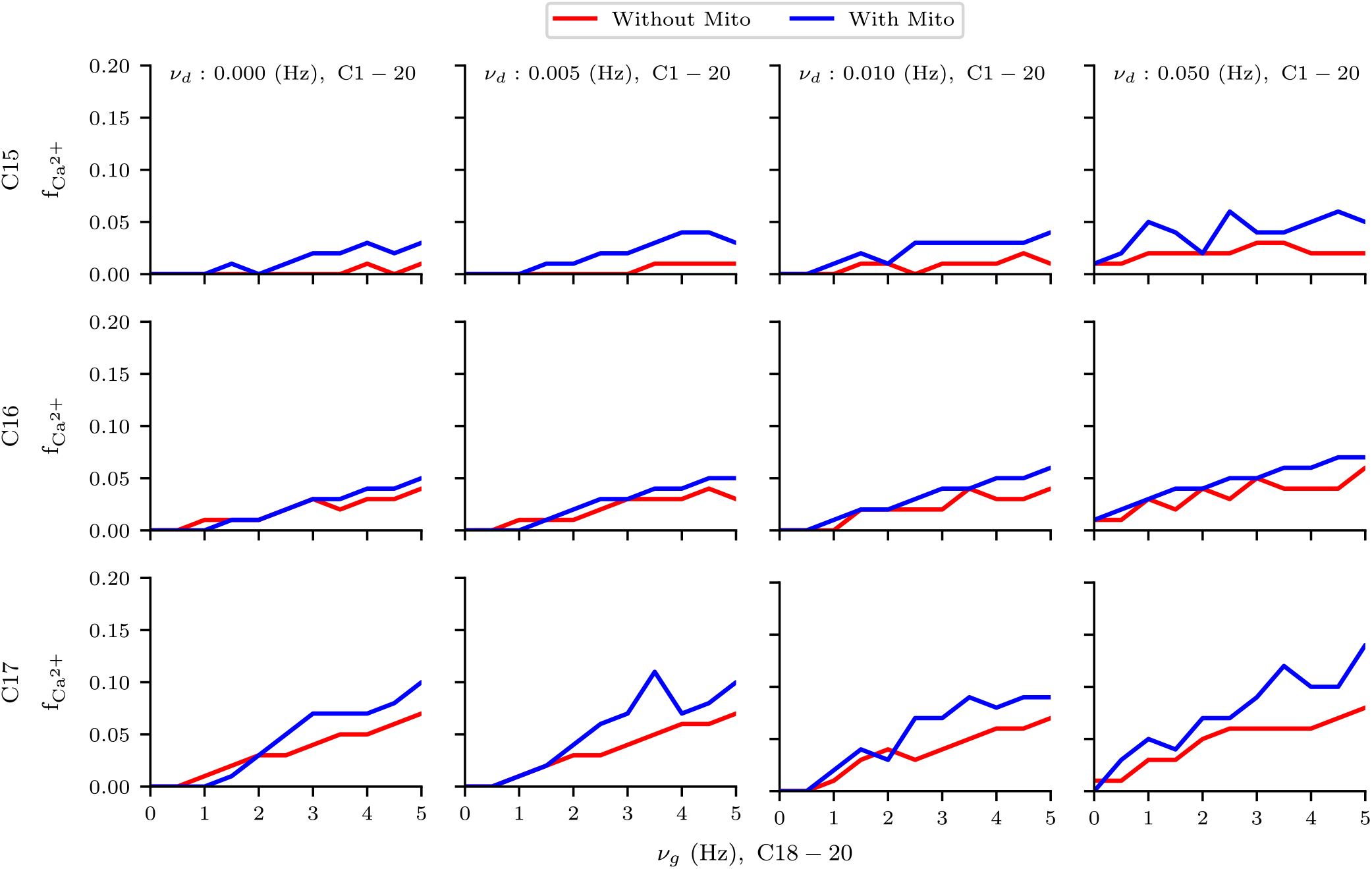
Mitochondria gates the transmission of Ca^2+^ signals in the linear astrocytic branch. The distal compartments are stimulated with glutamate (*v*_*g*_ from 0 to 5 Hz) while the whole cell is stimulated with dopamine (*v*_*d*_ = 0, 0.005, 0.01 or 0.05 Hz). **A** Schematic representation of the model morphology with the location of both inputs. **B** Propagated distance. **D** Difference in propagated distance between the models with and without mitochondria.

**FIGURE 8.**
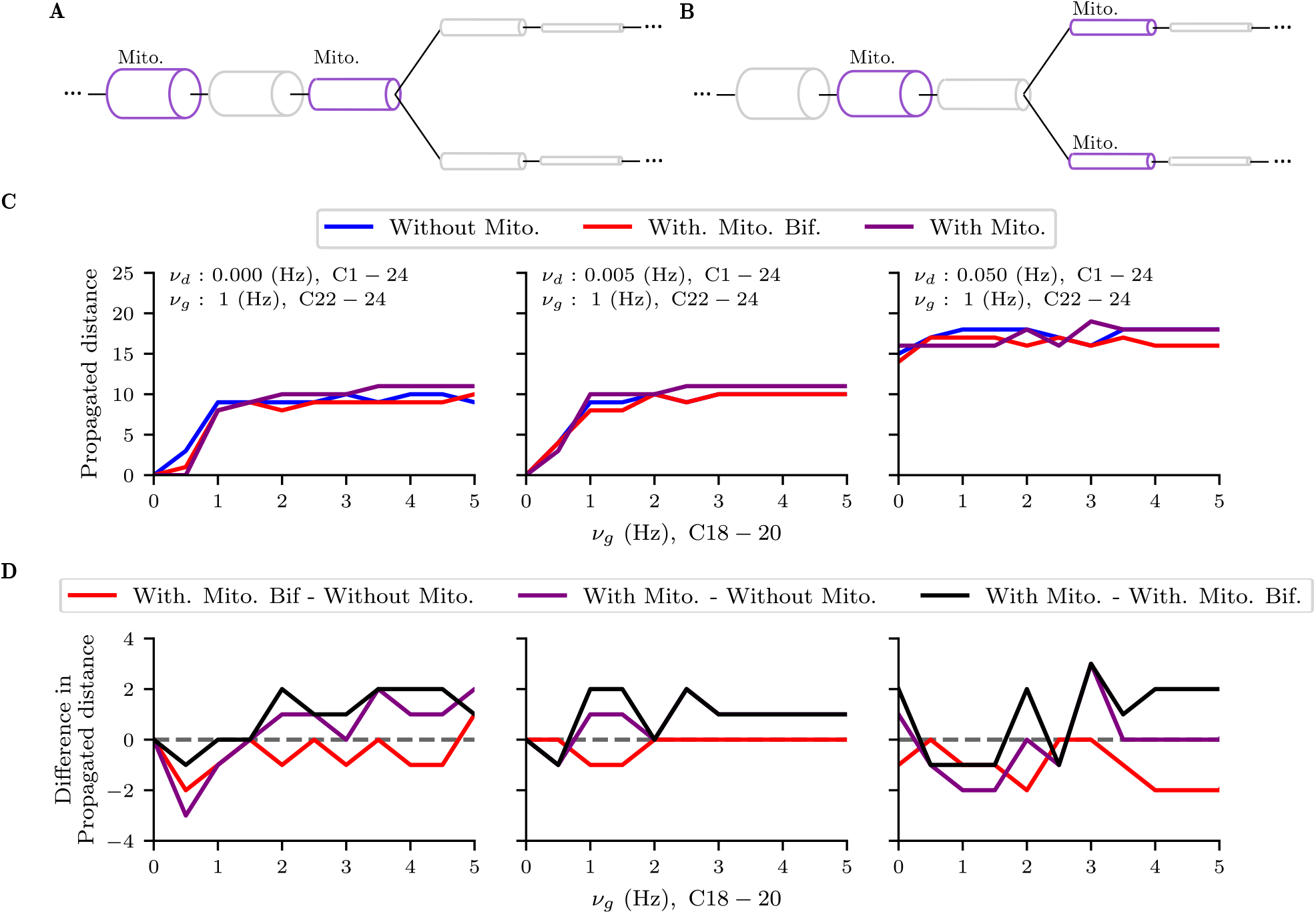
Mitochondria gates the transmission of Ca^2+^ signals from secondary branchlets. One branchlet is stimulated with fixed frequency glutamatergic input (*v*_*g*_ = 1 Hz), while the entire astrocyte is stimulated with dopamine (*v*_*d*_ = 0, 0.005 or 0.05 Hz), and the other branchlet with glutamate (*v*_*d*_ from 0 to 5 Hz.) Schematic representation of the model morphology with (**A**) and without (**B**) mitochondria at the bifurcation point. **C** Propagated distance. **D** Difference in propagated distance between the models without mitochondria, the model with mitochondria at the bifurcation, and the model without mitochondria at the bifurcation.

These results indicate that mitochondria have a bimodal effect depending on the strength of glutamatergic input and the presence of dopaminergic transmission. For low frequency glutamatergic input, mitochondria gate the transmission of Ca^2+^ signals from the distal compartments, reducing both the propagated distance and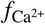. However, for stronger glutamatergic input and/or with dopaminergic stimulation, mitochondria increase the distance Ca^2+^ signals propagate from the terminal regions and the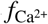.

### 3.3 Mitochondria gates the transmission of Ca^2+^ from terminal regions

Finally, since hierarchical branching of astrocytes processes influence how they interact ^15^ and as we detected that mitochondria gate the propagation of response from the distal regions, we further tested if these organelle could influence how secondary branchlets interact. In this test we used a bifurcated terminal morphology, composed of a single process emanating from the soma that bifurcates close to the terminal regions (Fig. 8**A** and **B**, and Tables 2 and 3), and studied if the presence of mitochondria at the branching point would change how the secondary branchlets interact. In one case we simulated a model with mitochondria at the branching point (named “With Mito. Bif.” condition, Fig. 8**A**), and the other without mitochondria (named “With Mito.”, Fig. 8**B**). The first secondary process is stimulated with different frequencies of glutamate in each trial (*v*_*g*_ from 0 to 5 Hz). The other branchlet is stimulated with a weak glutamatergic input (*v*_*g*_ = 1 Hz). The entire cell is stimulated with dopamine (*v*_*d*_ = 0, 0.005, 0.05 or 0.5 Hz). More details about the stimulation protocol are given in Text S1.

In absence of dopaminergic stimulation (*v*_*d*_ = 0) and low-frequency glutamatergic input (*v*_*g*_ < 1.5 Hz), mitochondria prevented the propagation of Ca^2+^ signals from the terminal regions in both models compared to the case without mitochondria (Fig. 8**C**). For any*v*_*g*_, mitochondria at the bifurcation point prevented the propagation of Ca^2+^ signals compared to the model without mitochondria at the bifurcation point (Fig. 8**C**). Interestingly, taking the difference between every model combination, the model with mitochondria at the bifurcation point reduced the propagated distance for any*v*_*g*_ compared to the model without mitochondria (Fig. 8**D**). In contrast, in the model without mitochondria at the bifurcation, mitochondria enhance the response in a similar manner to the unipolar morphology with mitochondria (Fig. 6**B**). Comparing the model with minus the model without mitochondria at the bifurcation, we observe that the presence of mitochondria at the bifurcation reduce the propagation distance. With low dopaminergic stimulation frequency (*v*_*d*_ = 0.005 Hz), the model with mitochondria at the bifurcation still show lower propagation distance compared to both the model without mitochondria and the model without mitochondria at the bifurcation (Fig. 8**C** and **D**). The model without the mitochondria at the bifurcation have similar response to the unipolar model with mitochondria (Fig. 6**B**, Fig. 8**C** and **D**). Even increasing the dopaminergic stimulation frequency, the propagation distance of the model with the mitochondria at the bifurcation is lower than the model without the mitochondria at the bifurcation and the model without any mitochondria (Fig. 7**C** and **D**).

Similar effect is observed for the 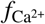. Here we focused the analysis of the response of compartments C16 (the bifurcation point), C17 and C21 (the adjacent compartments). Without dopaminergic stimulation, the response of compartments C17 and C21 in the model with mitochondria at the bifurcation point is similar to the model without any mitochondria (Fig. 9). In compartment C16, the 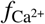is higher in the model with mitochondria at the bifurcation than the model without any mitochondria for stronger glutamatergic input (*v*_*g*_ > 3 Hz). With dopaminergic stimulation, the 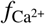of compartments C16, C17 and C21 is slightly higher than the model without any mitochondria (Fig. 9). Again, dopaminergic stimulation shifts the 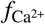curve to the left, so lower*v*_*g*_ already trigger higher 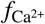in the both models with mitochondria compared to the model without mitochondria. However, in any case, the model without mitochondria at the bifurcation show higher 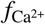than the model with the mitochondria at the bifurcation and the model without mitochondria, with a similar response profile to the unipolar morphology (Fig. 7 and Fig. 9). Together, these results show that the mitochondria at the bifurcation point gate the transmission of Ca^2+^ from the two secondary branchlets and prevent their communication, even for higher synaptic input at the first branchlet, in contrast with the unipolar morphology and with our previous study using a similar bifurcated terminal morphology ^15^.

**FIGURE 9.**
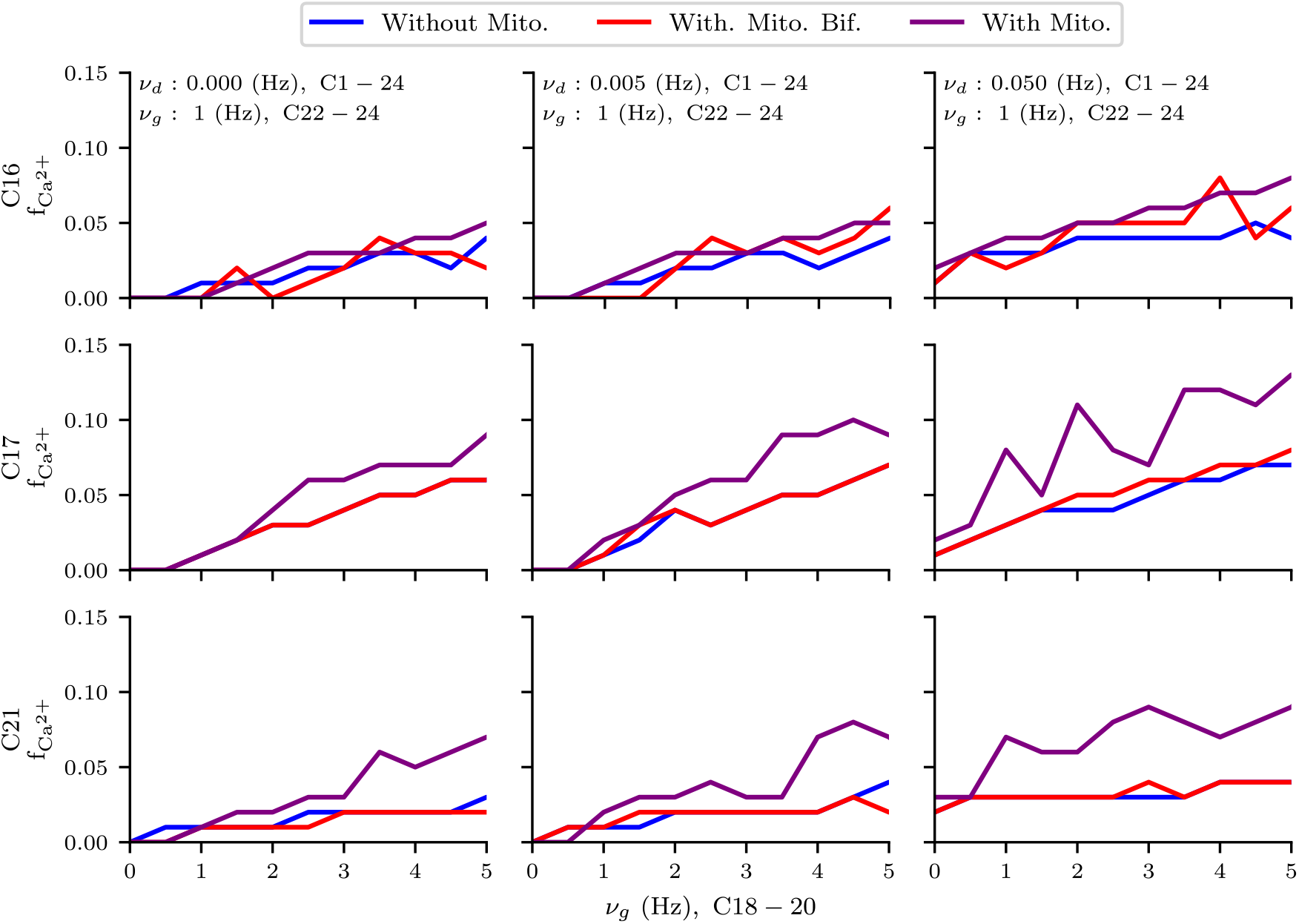
Mitochondria in the bifurcation point prevent the compartment response to Ca^2+^ signals from the secondary branchlets. All compartments are stimulated with dopamine (*v*_*d*_ = 0, 0.005 or 0.05 Hz), the second branchlet with glutamate (*v*_*g*_ = 1 Hz) and the first secondary branchlet with different frequencies of glutamatergic input (*v*_*g*_ from 0 to 5 Hz).

## 4 DISCUSSION

Here we present a computational model of astrocytes that include the MCU and mPTP mitochondrial mechanisms regulating intracellular Ca^2+^. We show that intramitochondrial Ca^2+^ follows closely the cytosol Ca^2+^ oscillation, and by uptaking Ca^2+^, they prevent the degradation of IP_3_, enhancing the response of the astrocyte and increasing the propagation distance in a context-dependent manner. For low glutamatergic input and in absence of dopaminergic input, these organelles reduce the frequency of Ca^2+^ oscillations and the propagation distance. In contrast, with dopaminergic stimulation or stronger glutamatergic input, mitochondria increase both the frequency of Ca^2+^ oscillations and the propagation distance. Mitochondria then gates the transmission of Ca^2+^ signals from the distal regions depending on the contextual information that an astrocyte receives.

Mitochondria are one of the major contributors to Ca^2+^ dynamics in astrocytes, uptaking Ca^2+^ from the cytosol through the MCU and releasing through mPTP ^2,3,53^. These mechanisms are involved in the regulation of synaptic plasticity in the hippocampus ^3^ and the generation of spontaneous Ca^2+^ microdomains in the mouse visual cortex ^2^. These mechanisms were also implemented in the present computational model. Here, mitochondria reduced the amplitude of the Ca^2+^ signals in the compartments with these organelles, but for strong glutamatergic input and with dopaminergic input increased the frequency of Ca^2+^ oscillations in every compartment. As the simulations suggest, mitochondria enhance the astrocyte response by transiently reducing the intracellular Ca^2+^ and then preventing the Ca^2+^-dependent IP_3_ degradation by the IP_3_-3K. These results are consistent with the subcellular location of mitochondria, since they are closely apposed to the ER, Ca^2+^ released from the ER can be readily uptake by mitochondria ^2,24^. The points an astrocyte is stimulated can be seen as “sources” of IP_3_ and non-stimulated compartments as “sinks” ^56,57,58^. By preventing the IP^3^ degradation, the mitochondria enable the accumulation of IP_3_ in the non-stimulated compartments and reduce the “sink” effect, enhancing the distance Ca^2+^ signals propagate. With dopaminergic stimulation, all compartments can be seen as IP_3_ sources, reducing the effect of the mitochondria over the IP_3_ accumulation. However, with weak glutamatergic input, the mitochondria already uptake the small amount of Ca^2+^ released, then preventing the astrocyte response. So, the present computational model predicts that Ca^2+^ signals detected closer to mitochondria should have lower amplitudes and that the propagation of Ca^2+^ signals and the size of the events should be associated with strong synaptic inputs or the release of a neuromodulator, such as dopamine and noradrenaline. Inhibition of mitochondria Ca^2+^ uptake and release, should reduce that propagation and the size of events when the astrocyte is subjected to strong synaptic input, but enhanced for weak glutamatergic stimulation.

Mitochondria also have a bimodal effect over the distance Ca^2+^ signals propagate from the terminal regions. While these organelles increase the propagation distance for high frequency glutamatergic input, they inhibit the propagation for low frequency stimulation. However, dopaminergic stimulation enables that even weaker glutamatergic inputs triggers Ca^2+^ signals that propagate from the distal compartments, that is, dopamine shifts the frequency threshold needed for glutamatergic stimulation to trigger a response that propagates. So, astrocytic mitochondria gate the propagation of Ca^2+^ signals from the distal compartments, preventing that weak glutamatergic input trigger a response that could propagate away the point of stimulation and prevent the transmission of Ca^2+^ signals from branchlets. Similarly, using a bifurcated morphology, we compared the propagation distance and the frequency of Ca^2+^ oscillation for the model with mitochondria at the bifurcation point and the model without mitochondria at the bifurcation. With this simulations we showed that the presence of mitochondria at the bifurcation prevented the propagation of Ca^2+^ signals from the distal regions and also inhibited the response of the neighboring compartments. So, similar to the unipolar morphology, mitochondria at bifurcation points gate the transmission of Ca^2+^ signals from the branchlets and inhibit the response in these regions.

Taking together, these results suggest that mitochondria prevent that lower synaptic activity, which could be associated with low demanding information processing, trigger global responses in astrocytes. In contrast, in the case with high neural activity, mitochondria further enhances the astrocyte responses. The point of transition from weak to strong stimulation is controlled by neuromodulators. Astrocytes have been linked to the codification of states, such as arousal, and states transition through neuromodulators ^59,10^. Since they are volume transmitters ^55,59,15^, these neuromodulators are particularly suited to trigger global events in astrocytes ^59^. So, in this sense, global events in astrocytes triggered by them could be associated with the codification of states and the mitochondria could then tune the astrocyte response to the network or task demand. Interestingly, the activity of dopaminergic neurons in the *substantia nigra* is also associated with the demand of cognitive task ^60^. Also, by preventing that weak synaptic inputs trigger global events, mitochondria could confine the astrocytic modulation to the synapses that are active. In contrast, in high demanding tasks, which would be reflected in higher neural activity and/or release of neuromodulators, the mitochondria enhance the astrocyte response, enabling the activation of the entire astrocyte and so the modulation of several synapses simultaneously, leading to broad modulation of neural networks and synchronization ^8^.

Ca^2+^ is also an important regulator of mitochondrial metabolism and, in particular, was associated with increase of ATP synthesis ^1^. Several regulators of intracellular Ca^2+^, as the SERCA pump, plasma membrane Ca^2+^ ATPase (PMCA), and the NCX, are energy dependent. Ca^2+^ uptake from the cytosol could be an important signaling mechanism of energy demand. Since astrocytes provide energy support for neurons, Ca^2+^ oscillations in astrocytes could also encode the neural network energy demand. Higher neural activity would induce higher frequency of Ca^2+^ oscillation, leading to higher mitochondrial metabolism.

Although not directly associated with the control of mitochondrial metabolism, a study showed that Ca^2+^ accumulation in astrocyte-like glia controls sleeping behavior in *Drosophila* ^61^. So, by preventing that Ca^2+^ signals propagate from terminal regions, mitochondria also prevent that weaker stimuli increase the mitochondrial metabolism in regions where it is not needed. However, during higher demanding tasks that require global modulation of neural networks and require more resources, Ca^2+^ responses can signal to the mitochondria the network energy demand and be further amplified by the mitochondria.

Studying mitochondria function is important for understanding different brain pathologies and psychiatry disorders. Alterations in astrocytes and mitochondria are associated with aging ^33^, Alzheimer’s and Parkinson’s diseases ^34,35^. In particular, reduced dopaminergic release, as observed in schizophrenia ^62^, could be associated with reduced global events in astrocytes ^15^, impacting the modulation of broad neural networks and network synchronization ^8,14,10^, and so be associated with the deficient codification of states and transition between them in schizophrenia ^63,64^. Interestingly, this disorder is also associated with altered metabolism and structure mitochondria in astrocytes ^36,37,38^. So, deficient coding of states and tuning of metabolic demand could be among the mechanisms by which dysfunction state transition emerges in schizophrenia and then a cause for the cognitive deficits in this disorder. So, mitochondria Ca^2+^ handling in astrocytes could be one of the potential targets to investigate treatments for schizophrenia.

## 5 CONCLUSIONS

Here we show that mitochondria influence Ca^2+^ signaling in astrocytes by preventing the Ca^2+^-dependent IP_3_ degradation. Mitochondria regulate Ca^2+^ signaling in astrocytes in a context-dependent manner that can be associated with tuning the astrocyte response to network demands. The present study is an important step in highlighting the mechanisms that control the Ca^2+^ response in astrocytes and could be one important link between neural and astrocytic activity with the regulation of metabolism in the brain.

## Abbreviations

ER: endoplasmic reticulum
MCU: mitochondrial calcium uniporter
mPTP: mitochondrial permeability transition pore.

## AUTHOR CONTRIBUTIONS

TOB and ACR contributed with the conceptualization, project administration, writing the original draft, review and editing. TOB developed the model implementation. ACR supervised the work.

